# SOX7 deficiency causes ventricular septal defects through its effects on endocardial-to-mesenchymal transition and the expression of *Wnt4* and *Bmp2*

**DOI:** 10.1101/2020.12.04.412288

**Authors:** Andrés Hernández-García, Sangbae Kim, Yumei Li, Bum Jun Kim, Hitisha P. Zaveri, Valerie K. Jordan, M. Cecilia Ljungberg, Rui Chen, Rainer B. Lanz, Daryl A. Scott

**Affiliations:** Department of Molecular and Human Genetics, Baylor College of Medicine, Houston, TX, USA; Department of Molecular Physiology and Biophysics, Baylor College of Medicine, Houston, TX, USA; Department of Pediatrics, Baylor College of Medicine, Houston, TX, USA; Jan and Dan Duncan Neurological Research Center at Texas Children’s Hospital, Houston, Texas, USA; Department of Molecular & Cellular Biology, Baylor College of Medicine, Houston, TX, USA; Quantitative and Computational Biosciences, Baylor College of Medicine, Houston, TX, USA

**Keywords:** SOX7, WNT4, BMP2, atrioventricular canal, endocardial-to-mesenchymal transition, ventricular septal defects

## Abstract

*SOX7* is located in a region on chromosome 8p23.1 that is recurrently deleted in individuals with septal defects. *Sox7*^−/−^ embryos die of heart failure around E11.5 due to defects in vascular remodeling. These embryos have hypocellular endocardial cushions with severely reduced numbers of mesenchymal cells. We also observed a ventricular septal defect in a rare *Sox7*^flox/−^;Tie2-Cre embryo that escaped early lethality. This led us to hypothesize that SOX7 plays a critical developmental role in the endocardium of the atrioventricular (AV) canal. We subsequently used AV explant studies to show that SOX7 deficiency leads to a severe reduction in endocardial-to-mesenchymal transition (EndMT). Since SOX7 is a transcription factor, we hypothesized that it functions in the endocardium by regulating the expression of EndMT-related genes. To identify these genes in an unbiased manner, we performed RNA-seq on pooled E9.5 hearts tubes harvested from *Sox7*^−/−^ embryos and their wild-type littermates. We found that *Wnt4* transcript levels were severely reduced, which we confirmed by RNA in situ hybridization. Previous studies have shown that WNT4 is expressed in the endocardium and promotes EndMT by acting in a paracrine manner to increase the expression of BMP2 in the myocardium. Consistent with these findings, we found that *Bmp2* transcript levels were diminished in *Sox7*^−/−^ embryonic hearts. We conclude that SOX7 promotes EndMT in the developing AV canal by modulating the expression of *Wnt4* and *Bmp2*. These data also provide additional evidence that haploinsufficiency of *SOX7* contributes to the congenital heart defects seen in individuals with recurrent 8p23.1 microdeletions.

**SUMMARY STATEMENT:** In the developing atrioventricular canal, SOX7 promotes endocardial-to-mesenchymal transition (EndMT) by positively regulating *Wnt4* and *Bmp2* expression. SOX7 deficiency leads to the development of hypocellular endothelial cushions and ventricular septal defects.

## INTRODUCTION

8p23.1 microdeletion syndrome is caused by a non-allelic homologous recombination mediated by low-copy repeats flanking an ~3.4 Mb region of DNA (Wat et al., 2009). Recurrent deletions of this region are associated with a variety of neurodevelopmental phenotypes, dysmorphic features a high incidence of congenital heart defects (CHD) and congenital diaphragmatic hernia (Devriendt et al., 1999; Lopez et al., 2006; Shimokawa et al., 2005; Slavotinek et al., 2005). Haploinsufficiency of *GATA4*, one of the genes located in this region, is sufficient to cause CHD (Garg et al., 2003; Okubo et al., 2004). However, published reports suggested that the spectrum of cardiac defects seen in individuals with 8p23.1 deletions is more severe than that seen in individuals with heterozygous pathogenic variants in *GATA4* (Wat et al., 2009). This led to the hypothesis that haploinsufficiency of other genes on chromosome 8p23.1 may also be contributing to the development of CHD.

*SOX7* is located with *GATA4* in the recurrently deleted region on chromosome 8p23.1 and has been hypothesized to contribute to the development of CDH associated with deletions of this region (Wat et al., 2009). Animal models provide support for the role of SOX7 in cardiovascular development. In Xenopus, injection of RNAs encoding *Sox7* leads to the nodal-dependent expression of markers of cardiogenesis in animal cap explants, while injection of morpholinos directed against *Sox7* lead to a partial inhibition of cardiogenesis in vivo (Zhang et al., 2005). In Zebrafish, simultaneous knockdown of *Sox7* and *Sox18* leads to a severe loss of the arterial identity of the presumptive aorta leading to dysmorphogenesis of the proximal aorta, the development of arteriovenous shunts, and a lack of circulation in the trunk and tail (Herpers et al., 2008; Pendeville et al., 2008). In mice, ablation of *Sox7* resulted in defects in vascular remodeling. Specifically, *Sox7*^−/−^ embryos die around E11.5 with dilated pericardial sacs and failure of yolk sac remodeling suggestive of cardiovascular failure (Wat et al., 2012). However, these studies have failed to demonstrate a clear association between decreased expression of SOX7 and the development of the types of CHD typically seen in individuals with 8p23.1 microdeletion syndrome.

Here we show that SOX7 deficiency in mice leads to the development of hypoplastic endocardial cushions and ventricular septal defects caused by a lack of endocardial-to-mesenchymal transformation (EndMT) in the developing atrioventricular (AV) canal. We go on to show that SOX7 modulates the expression of *Wnt4* in the endocardium and, secondarily, *Bmp2* in the myocardium; two genes previously shown to be required for EndMT in the AV canal. These data also provide additional evidence that haploinsufficiency of *SOX7* contributes to the congenital heart defects seen in individuals with recurrent 8p23.1 microdeletions.

## RESULTS

### SOX7 deficiency leads to disruption of ventricular septation

*Sox7*^−/−^ mouse embryos die around E11.5 (Wat et al., 2012), a time point that does allow evaluation of the development of the AV septum. In an effort overcome this limitation, we attempted to generate *Sox7*^flox/−^;Tie-2 Cre embryos in which one *Sox7* allele was ablated systemically and the other was selectively ablated in endothelial cells, including the endocardium, using a transgenic Tie2-Cre (Kisanuki et al., 2001) which does not affect the expression of the *Tie2* (previously known as *Tek*). Although the majority of these embryos died prior to E15.5, we were able to harvest two rare survivors at this time point. One of these embryos had normal cardiac anatomy, but the other had a ventricular septal defect not seen in *Sox7*^flox/−^ control littermates (Fig. 1A, B).

**Figure 1.**
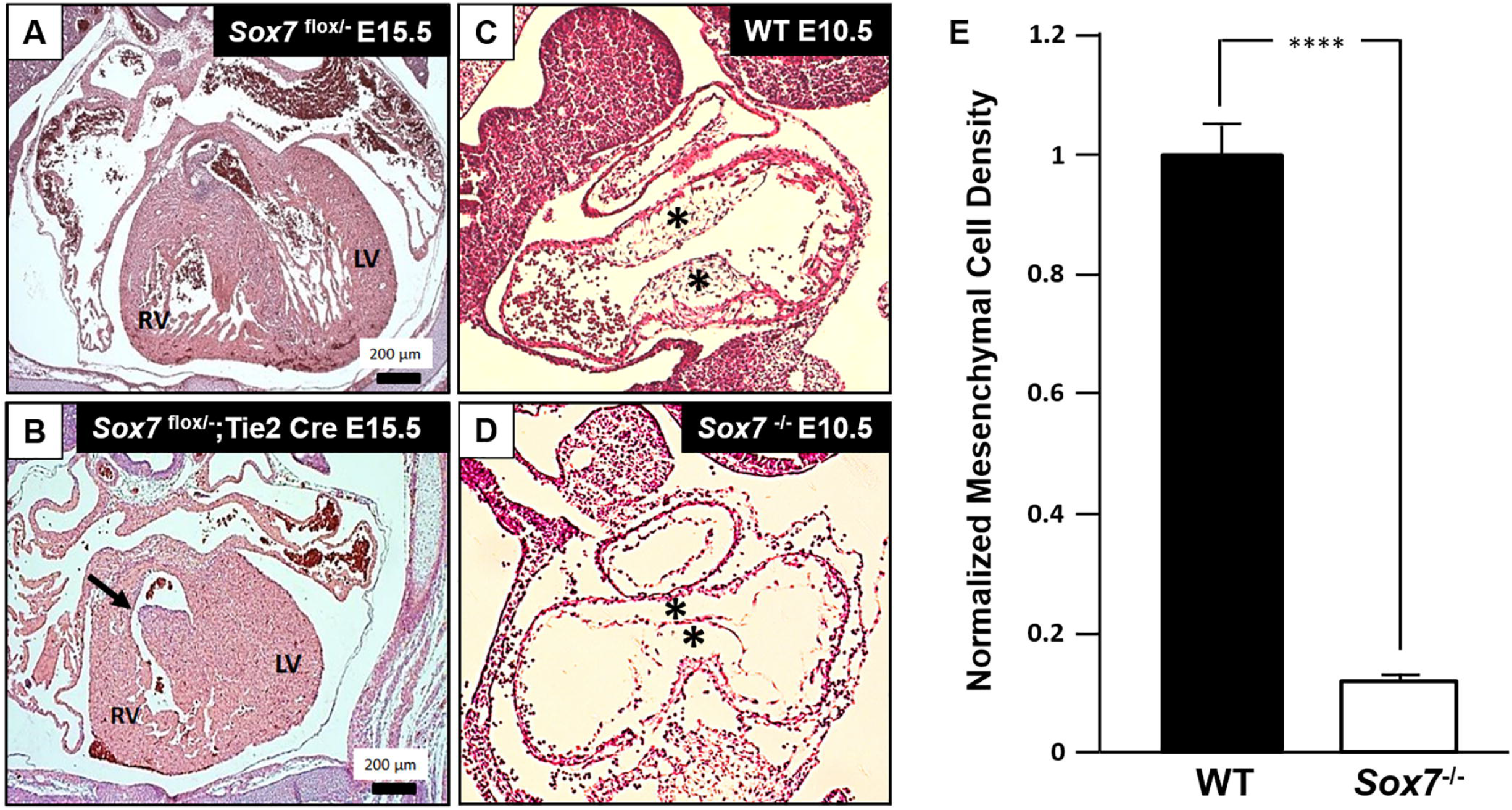
SOX7 deficiency causes ventricular septal defects and hypocellular AV cushions. A-B) One of two rare *Sox7*^flox/−^;Tie-2 Cre embryos harvested at E15.5 had a ventricular septal defect (black arrow in B). C-D) At E10.5, the AV endocardial cushions of wild-type embryos are filled with mesenchymal cells (*). In contrast, the AV endocardial cushions of *Sox7*^−/−^ embryos harvested at E10.5 are hypocellular (*). E) The normalized mesenchymal cell densities of wild-type and *Sox7*^−/−^ embryos were calculated based on 6-8 sagittal sections through the heart obtained from each of three embryos of each genotype. *Sox7*^−/−^ embryos have significantly reduced mesenchymal cell density in their AV canals compared to wild-type embryos (P < 0.0001). Error bars represent the standard error of the mean. LV= left ventricle, RV = right ventricle.

### SOX7 deficiency leads to hypocellular AV endocardial cushions

Septal defects are the most common form of CHD associated with 8p23.1 microdeletions (Wat et al., 2009). Since the mesenchymal cells of the AV endocardial cushions will ultimately form the AV septum and its associated valves (Anderson et al., 2003), we looked for difference in the development of the endocardial cushions in E10.5 Sox7^−/−^ embryos when compared to their wild-type littermates. Examination of sagittal sections through the AV endocardial cushions of *Sox7*^−/−^ embryos revealed normal separation of the endocardium from the underlying myocardium. Furthermore, the size of the AV endocardial cushions in *Sox7*^−/−^ embryos were comparable to those of their wild-type littermates (Fig. 1C-E). In contrast, the AV endocardial cushions of *Sox7*^−/−^ embryos were had severely reduced numbers of mesenchymal cells when compared to those of their wild-type littermates.

### SOX7 deficiency leads to a decrease in EndMT

During septal development, a subset of endocardial cells in the AV canal undergo EndMT to form mesenchymal cells, which migrate into the cardiac jelly between the endocardium and the myocardium (Armstrong and Bischoff, 2004; Markwald et al., 1981). Hence, decreased levels of EndMT can lead to development of hypocellular AV endocardial cushions. To determine if a defect in EndMT could underlie the hypocellularity of the AV endocardial cushions seen in *Sox7*^−/−^ embryos, we performed collagen gel AV canal explant studies. Briefly, the AV canals of *Sox7*^−/−^ embryos and their wild-type littermates were harvested at E9.5, and the number of migrating mesenchymal cells emanating from the explants were counted after 48 hours of collagen gel culture. These studies revealed a significantly reduced numbers of migrating mesenchymal cells associated with *Sox7*^−/−^ explants compared to wild type explants (P <0.0001, Fig 2). This suggest that a defect in EndMT is likely to be a major contributor to the development of the hypocellular AV endocardial cushions caused by ablation of *Sox7*.

**Figure 2.**
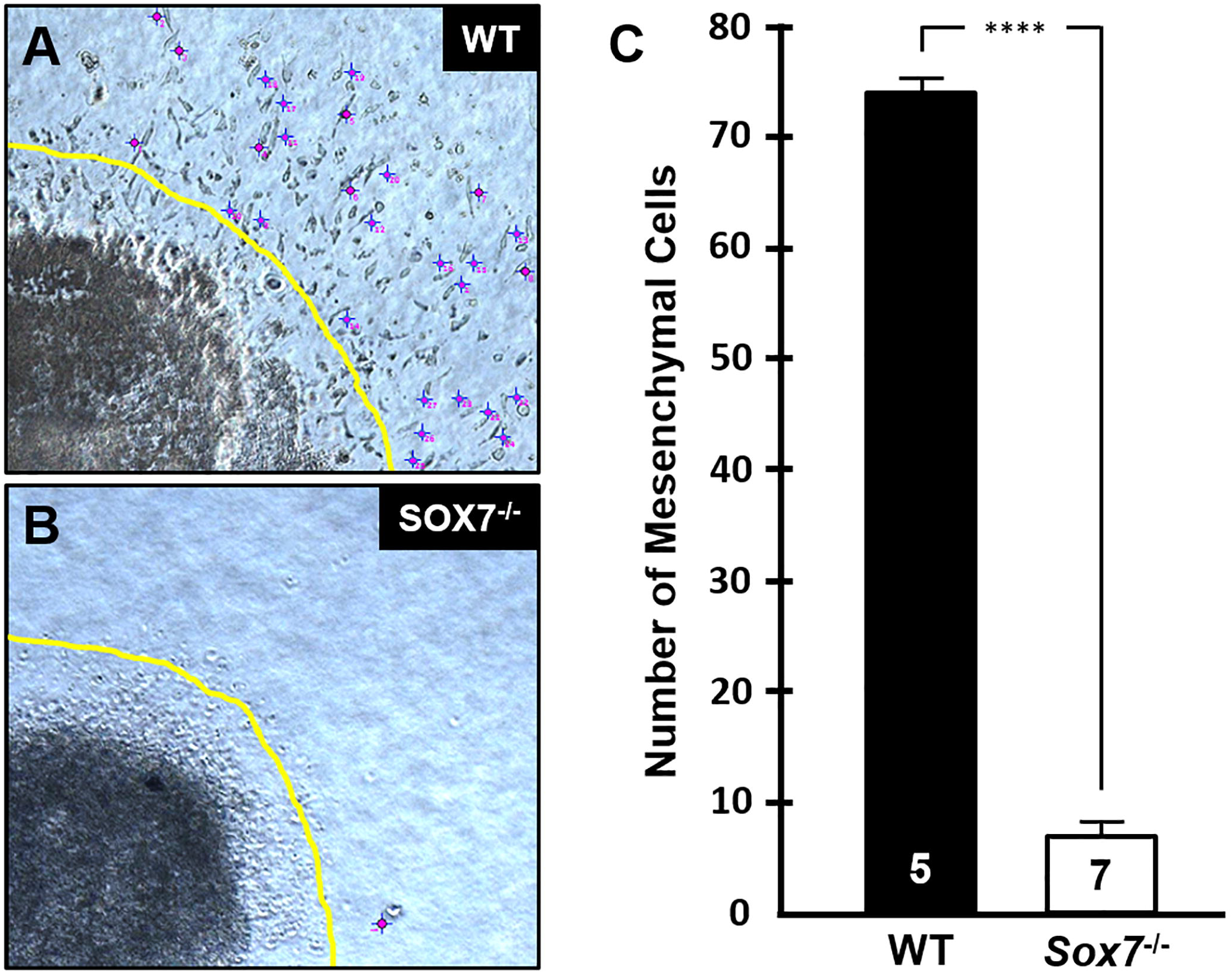
Ablation of *Sox7* leads to a decreased in EndMT. A) Collagen gel AV canal explants from E9.5 wild-type embryos generated a large number of migrating mesenchymal cells (marked by purple crosses). B-C) In contrast, collagen gel AV canal explants from <u>*Sox7*^−/−^</u> embryos generated a significantly reduced number of migrating mesenchymal cells (P < 0.0001) suggesting that SOX7 deficiency causes a severe defect in EndMT. Figures and are representative images from explants obtained from five wild-type and seven *Sox7*^−/−^ embryos generated in crosses of *Sox7*^+/−^ mice on a C57BL/6 background. Error bars represent the standard error of the mean.

### RNA-seq analysis reveals alterations in genes associated with epithelial-to-mesenchymal transition with down-regulation of *Wnt4* and *Bmp2*

Since *Sox7* encodes a DNA binding transcription factor, it is reasonable to assume that it regulates the transcription of key genes during heart development (Takash et al., 2001; Taniguchi et al., 1999). To identify Sox7 target genes in an unbiased manner, we compared the RNA-seq transcriptomes of pooled heart tubes from *Sox7*^−/−^ embryos harvested at E9.5 and those to those of their wild-type littermates. Using the Bioconductor DESeq2 package, we found 722 differentially expressed genes (p-value < 0.001 and absolute fold change > 2; Supplemental Table 1). We then queried the Molecular Signature Database (MSigDB) with Gene Set Enrichment Analysis software (Mootha et al., 2003; Subramanian et al., 2005) to determine whether the *Sox7*^−/−^ differentially expressed genes show statistically significant, concordant differences between biological states. Figure 3A shows ranked overlaps with the MSigDB Hallmark gene sets, which summarize and represent well-defined phenotypes. Ingenuity Canonical Pathway analysis also revealed enrichments for the canonical pathways ‘Glycolysis’, ‘Nitric Oxide Signaling in the Cardiovascular System’, and ‘Regulation of the Epithelial-Mesenchymal Transition Pathway’. Overlapping genes for each Hallmark gene set are shown in Supplemental Table 2. ‘Cardiovascular System Development and Function’ was the top enrichment (p-value 3.42e-16) for disease and function annotations.

**Figure 3.**
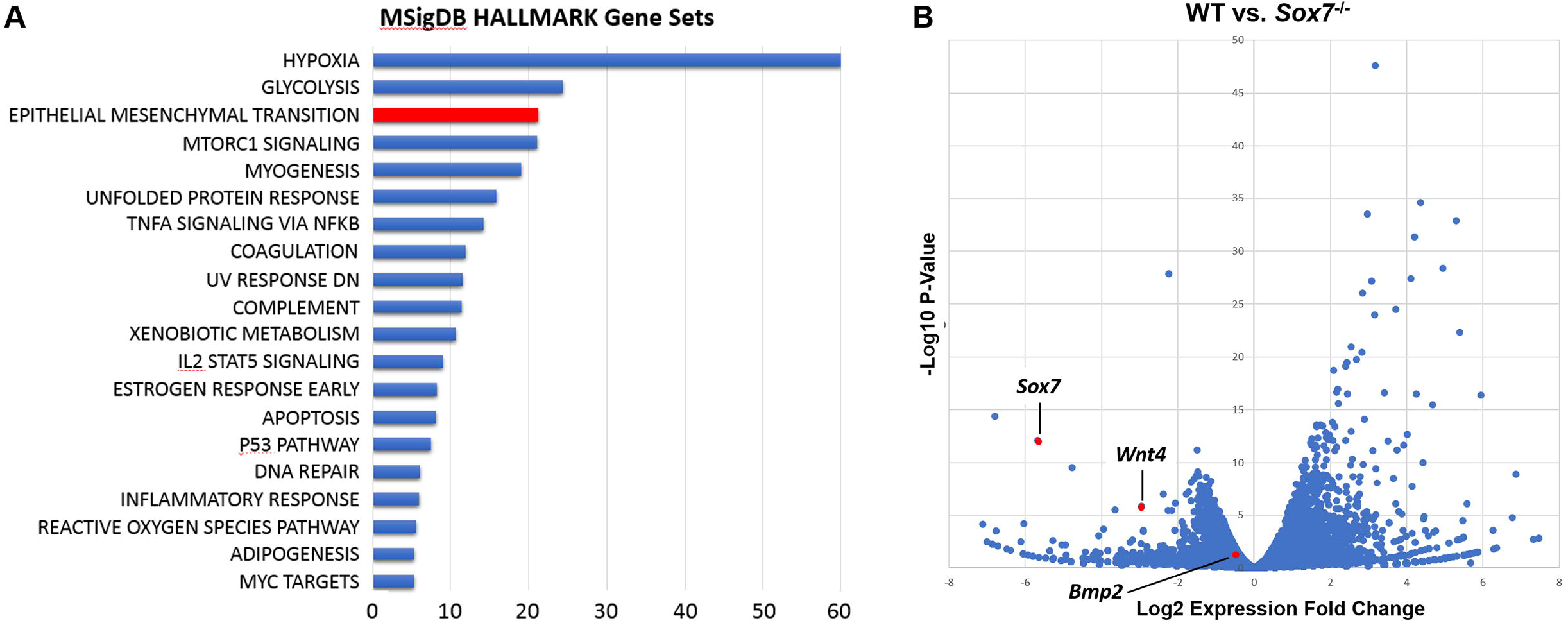
RNA-seq analyses of E9.5 wild-type and *Sox7*^+/−^ heart tubes reveals perturbations in genes involved in epithelial to mesenchymal transition and a decrease in *Wnt4* and *Bmp2* transcripts. A) MSigDB Hallmark gene set enrichments for 722 genes that were differentially expressed with p-value < 0.001 and absolute fold change > 2. B) Volcano scatter plot representation of differentially expressed genes. We found significantly decreased levels of *Wnt4* expression levels were significantly decreased. To a lesser extent, the expression level of WNT4’s downstream target, *Bmp2*, was also decreased.

Among the differentially expressed genes with known roles in EndMT, we noted that *Wnt4* expression was downregulated (Figure 3B). WNT4 is expressed in the endocardium and promotes EndMT by acting in a paracrine manner to increase the expression of BMP2 in the myocardium (Wang et al., 2013). Consistent with these findings, we found that *Bmp2* transcript levels were also diminished in *Sox7*^−/−^ embryonic hearts, although to a lesser degree (Figure 3B).

### *Wnt4* and *Bmp2* transcript levels are decreased in the atrioventricular canals of *Sox7*^−/−^ mice

To confirm the downregulation of *Wnt4* and *Bmp2*, we compared their transcript level in the AV canals of *Sox7*^−/−^ and wild-type embryos at E9.5 by RNA in situ hybridization (RNA-ISH). Both *Wnt4* and *Bmp2* transcript levels were reduced in AV canal sections obtained from E9.5 *Sox7*^−/−^ embryos compared to those obtained from their wild-type littermates (Figure 4).

**Figure 4.**
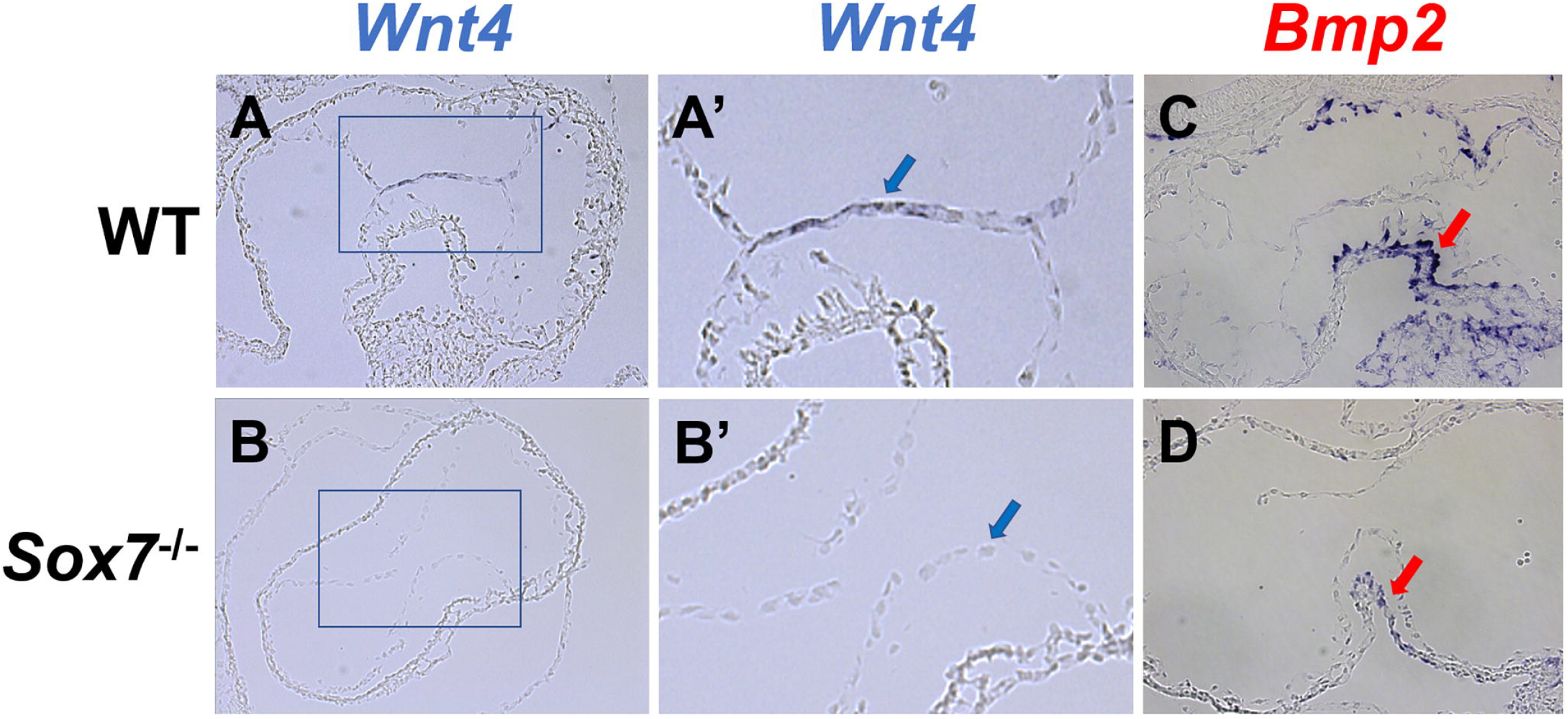
SOX7 deficiency leads to decreased expression of *Wnt4* and *Bmp2* in the developing AV canal at E9.5. A, A’) RNA-ISH performed on sections obtained from E9.5 wild-type embryos demonstrates that *Wnt4* is expressed in the endocardium covering the AV endocardial cushions (blue arrow in A’). B, B’) *Sox7*^−/−^ embryos show reduced levels of *Wnt4* transcripts in the AV endocardium when compared to wild-type embryos (blue arrow in B’). C, D) *Bmp2* transcripts are detected in the myocardium of the AV canal with reduced levels being seen in *Sox7*^−/−^ embryos (red arrow in D) compared to wild-type controls (red arrow in C). Figures are representative images from RNA-ISH studies performed on sections obtained from four embryos of each genotype that were generated in crosses of *Sox7*^+/−^ mice on a C57BL/6 background.

## DISCUSSION

CHD is present in over 1% of all newborns and are the leading cause of birth-defect-related deaths (Gilboa et al., 2010; Jenkins et al., 2007; Pierpont et al., 2007a). The majority of CHD cases in the general population are thought to be caused by multiple genetic and environmental factors interacting in a multifactorial mode of inheritance (Pierpont et al., 2007b). These types of interactions can be difficult to study in the general population due to their complexity. One means of reducing this complexity is to study microdeletions in which the major CDH candidate genes are limited based on their location within a specific region.

8p23.1 microdeletion syndrome is associated with a high incidence of CDH with septal defects being particularly common (Wat et al., 2009). Although haploinsufficiency of *GATA4* is sufficient to cause CHD (Garg et al., 2003; Okubo et al., 2004; Pehlivan et al., 1999; Reamon-Buettner and Borlak, 2005; Sarkozy et al., 2005), clinical data suggests that haploinsufficiency of at least one additional gene in this region promotes the development of CDH. Haploinsufficiency of *SOX7* has been previously hypothesized to contribute to the development of CDH associated with 8p23.1 microdeletions (Wat et al., 2009). Here we use mouse models to elucidate the morphogenetic and molecular mechanisms by which SOX7 deficiency causes septal defects in mice.

### SOX7 deficient embryos have hypocellular AV cushions and develop ventricular septal defects due to decreased levels of EndMT

### SOX7 regulates EndMT through its effects on *Wnt4* and *Bmp2* transcription

Since SOX7 is an endocardially expressed DNA binding transcription factor (Takash et al., 2001; Taniguchi et al., 1999; Wat et al., 2012), and its deficiency causes a severe defect in EndMT (Figure 2), we hypothesized that it functions in a cell autonomous fashion to regulate the transcription of genes that regulate EndMT. RNA-seq performed on E9.5 heart tubes (Figure 3B) and RNA-ISH studies (Figure 4) revealed decreased expression of *Wnt4* and, to a lesser extent, *Bmp2*. During the development of the AV endocardial cushions, WNT4 is expressed in the endocardium and acts as a paracrine factor to upregulate *Bmp2* expression in the adjacent myocardium (Wang et al., 2013). In turn, BMP2 functions to induce EndMT in the AV canal (Ma et al., 2005). In humans, homozygous loss-of-function variants in *WNT4* have been reported to cause autosomal recessive 46,XX sex reversal with dysgenesis of kidney, adrenals, and lungs (SERKAL syndrome) whose features also include ventricular septal defects (Mandel et al., 2008). Similarly, heterozygous variants in *BMP2* cause short stature, facial dysmorphism, and skeletal anomalies with or without cardiac anomalies (SSFSC syndrome) that include ventricular septal defects (Tan et al., 2017). We conclude that SOX7 modulates EndMT in the developing AV canal by modulating the expression of *Wnt4* and its downstream target gene, *Bmp2* (Figure 5).

**Figure 5:**
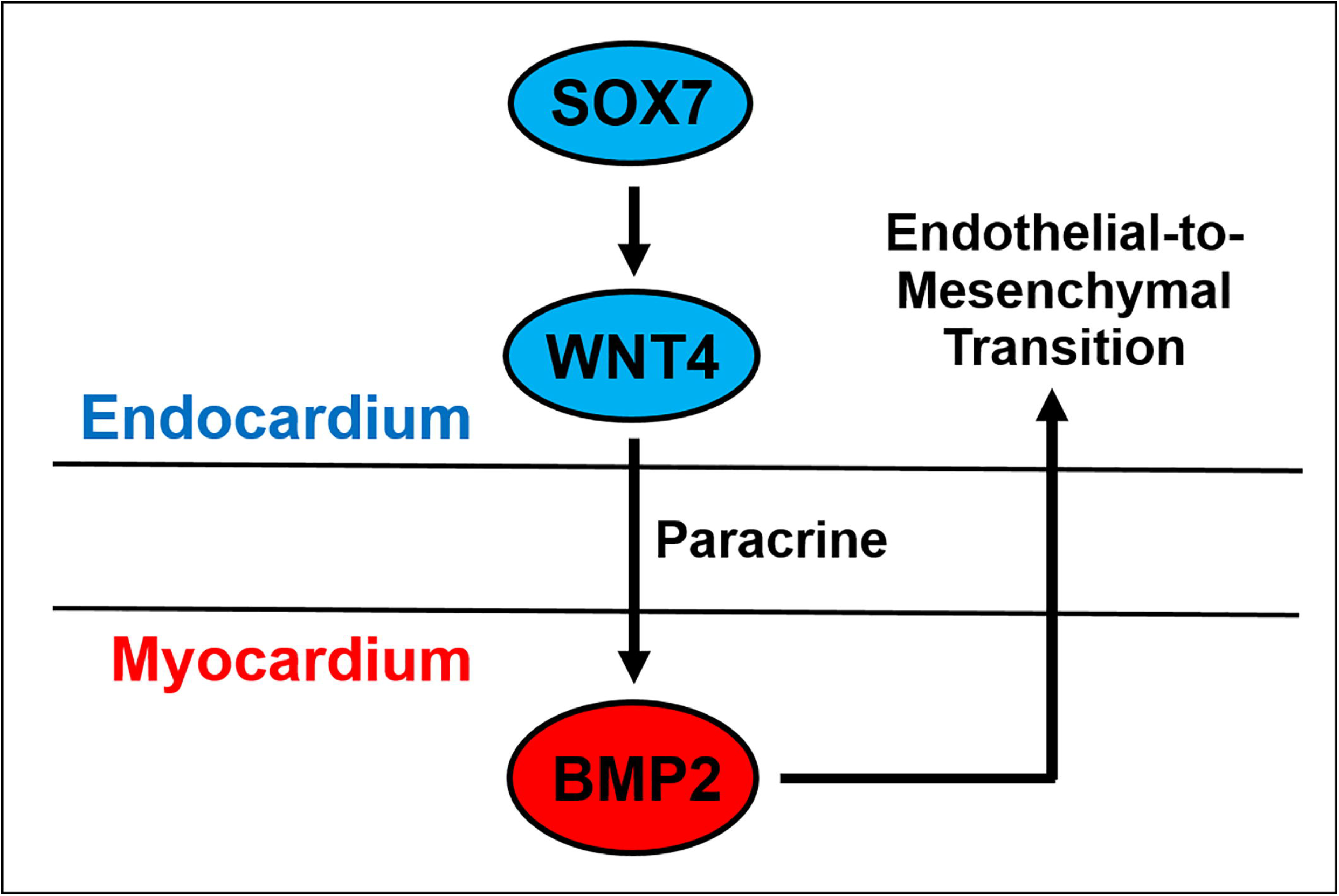
The role of SOX7 in the development of the AV endocardial cushions. SOX7 functions in the endocardium to positively regulate the expression of *Wnt4*. WNT4 acts as a paracrine factor to upregulate *Bmp2* expression in the myocardium (Wang et al., 2013). BMP2 then functions to induce EndMT in the AV canal (Ma et al., 2005). In *Sox*7^+/−^ embryos, a defect in EndMT leads to the development of hypocellular AV endocardial cushions and ventricular septal defects.

## MATERIALS AND METHODS

### Mouse models

Generation of mice bearing *Sox7* conditional (flox) and null alleles were described previously (Wat et al., 2012). Briefly, the second exon of *Sox7*, which encodes half of SOX7′s HMG-box DNA binding domain and the entire SOX7 activation domain was flanked by loxP sites to generate the flox allele. In the presence of Cre, the second exon and the 3’ untranslated region are excised generating a null allele.

All mice in our study were maintained on a C57BL/6 background. All experiments using mouse models were conducted in accordance with the recommendations in the Guide for the Care and Use of Laboratory Animals of the National Institutes of Health (NIH). The associated protocols were approved by the Institutional Animal Care and Use Committee of Baylor College of Medicine (Animal Welfare Assurance #A3832-01).

### Histology

For standard histology, embryos were fixed in buffered Formalde-Fresh Solution (Fisher) or 4% buffered paraformaldehyde (PFA). After fixation the specimens were washed in PBS, dehydrated in ethanol, embedded in paraffin and sectioned at 10 μm. Series of sections were then stained with hematoxylin and eosin (H&E).

### Collagen gel explant assays

Collagen gel explant assays were performed as described by Xiong et al. (Xiong et al., 2012). Briefly, collagen gels were prepared from a type I rat-tail collagen stock at ~4mg/ml (Corning). The solution was diluted with 1x PBS and buffered with 1 N NaOH. Collagen gels were soaked with culture medium containing OptiMEM-I (Gibco), 1% FBS (Thermo Fisher Scientific), 100 U/ml penicillin (Thermo Fisher Scientific) and 100 μg/ml streptomycin (Thermo Fisher Scientific) overnight at 37°C, and were allowed to solidify inside a 5% CO2 incubator at 37°C. The culture medium was removed from the gels prior to the dissection of the AV canals.

E9.5 embryos with 20-25 pairs of somites, were harvested in cold PBS. After elimination of the pericardial sac, the AV canals were dissected and placed with the endocardium facing downwards on the surface of a collagen gel. Explants were incubated in 5% CO2 at 37°C. Images were acquired at 48 hours of culture using an Axiovert 40 CFL inverted microscope equipped with an AxioCam digital camera and imaging system (Carl Zeiss Microscopy).

Migratory mesenchymal cells (hypertrophied, elongated spindle-shaped morphology, and growing out from the explant marginal area) were counted from explants obtained from five wild-type and seven *Sox7*^−/−^ embryos. An unpaired two-tailed Student’s t-test was used to compare the numbers of migratory mesenchymal cells generated from explants of different genotypes.

### E9.5 heart tube RNA isolation and RNA-seq analysis

Heart tubes were isolated from *Sox7*^−/−^ embryos and their C57BL/6 wild-type littermates at E9.5. Briefly, heart tubes were micro dissected at the distal part of the outflow tract before it opens within the region of the aortic sac. Next, a lower slice was performed at the end of the common atrial chamber to obtain a heart segment including the endocardial cushions and the AV canal. Cephalic and tail poles were saved for genotyping. RNA was obtained from pools of approximately 30-50 heart tubes of each genotype using an RNeasy Plus Micro Kit (QIAGEN) according to manufacturer’s instructions. RNA from two pooled *Sox7*^−/−^ samples and one pooled wild-type sample were used for RNA-seq analyses.

Bulk RNA-Seq were performed at the Department of Molecular and Human Genetics Functional Genomics Core at Baylor College of Medicine. RNAseq libraries ware made using a KAPA stranded mRNA-seq kit (KK8420). Briefly, poly-A RNA was purified from total RNA using Oligo-dT beads, fragmented to a small size, after which first strand cDNA was synthesized. Second strand cDNA was synthesized and marked with dUTP. The resultant cDNA was used for end repair, A-tailing and adaptor ligation. Finally, libraries were amplified for sequencing using the Novaseq platform (Illumina). The strand marked with dUTP was not amplified, allowing strand-specific sequencing.

### RNA in situ hybridization

Embryos (E9.5) were collected, washed in cold PBS, and fixed overnight in 4% PFA. After washing, the embryos were cryoprotected sequentially in PBS buffered 15% and 30% sucrose solutions, embedded in OCT (Tissue-Tek), and snap frozen. RNA in situ hybridization was performed on serial 10 and 12 μm thick sections cuts on a Leica CM3050 S cryostat. Sections were probed with digoxigenin-labelled sense and anti-sense mRNA probes. The *Wnt4* probe included a 800 bp sequence based on NM_009523.2 flanked by forward 5’- CAGCATCTCCGAAGAGGAGAC-3’ and reverse 5’-CTTTAGATGTCTTGTTGCACG-3’ sequences. The *Bmp2* probe included a 259 bp region sequence based on NM_007553.1 flanked by forward 5’- GATCTTCCGGGAACAGATACAG-3’ and reverse 5’- CACCTGGGTTCTCCTCTAAATG-3’ sequences as previously described in the Allen Brain Atlas (Lein et al., 2007). In situ hybridization was performed by the RNA In Situ Hybridization Core at Baylor College of Medicine using an automated robotic platform and a previously described protocol (White et al., 2014; Yaylaoglu et al., 2005). Images were acquired using a Leica DM4000 microscope equipped with a Leica DMC 2900 camera.

## ACKNOWLEDGEMENTS

RNA-ISH was performed in the RNA In Situ Hybridization Core at Baylor College of Medicine). RNA-seq was performed in the Department of Molecular and Human Genetics Functional Genomics Core at Baylor College of Medicine.

## DATA AVAILABILITY

Primary RNA-seq data will be made available in the Short Reach Archive.

## REFERENCES

Anderson, R. H., Webb, S., Brown, N. A., Lamers, W. and Moorman, A. (2003). Development of the heart: (2) Septation of the atriums and ventricles. Heart 89, 949–58.

Armstrong, E. J. and Bischoff, J. (2004). Heart valve development: endothelial cell signaling and differentiation. Circ. Res. 95, 459–70.

Combs, M. D. and Yutzey, K. E. (2009). Heart valve development: regulatory networks in development and disease. Circ. Res. 105, 408–21.

Devriendt, K., Matthijs, G., Van Dael, R., Gewillig, M., Eyskens, B., Hjalgrim, H., Dolmer, B., McGaughran, J., Brondum-Nielsen, K., Marynen, P. et al. (1999). Delineation of the critical deletion region for congenital heart defects, on chromosome 8p23.1. Am. J. Hum. Genet. 64, 1119–26.

Eisenberg, L. M. and Markwald, R. R. (1995). Molecular regulation of atrioventricular valvuloseptal morphogenesis. Circ. Res. 77, 1–6.

Garg, V., Kathiriya, I. S., Barnes, R., Schluterman, M. K., King, I. N., Butler, C. A., Rothrock, C. R., Eapen, R. S., Hirayama-Yamada, K., Joo, K. et al. (2003). GATA4 mutations cause human congenital heart defects and reveal an interaction with TBX5. Nature 424, 443–7.

Gilboa, S. M., Salemi, J. L., Nembhard, W. N., Fixler, D. E. and Correa, A. (2010). Mortality resulting from congenital heart disease among children and adults in the United States, 1999 to 2006. Circulation 122, 2254–63.

Herpers, R., van de Kamp, E., Duckers, H. J. and Schulte-Merker, S. (2008). Redundant roles for sox7 and sox18 in arteriovenous specification in zebrafish. Circ. Res. 102, 12–5.

Jenkins, K. J., Correa, A., Feinstein, J. A., Botto, L., Daniels, S. R., Elixson, M., Warnes, C. A., Webb, C. L. and Young, American Heart Association Council on Cardiovascular Disease in the Young (2007). Noninherited Risk Factors and Congenital Cardiovascular Defects: Current Knowledge: A Scientific Statement From the American Heart Association Council on Cardiovascular Disease in the Young: Endoresed by the American Academy of Pediatrics. Circulation 115, 2995–3014.

Kaneko, K., Li, X., Zhang, X., Lamberti, J. J., Jamieson, S. W. and Thistlethwaite, P. A. (2008). Endothelial expression of bone morphogenetic protein receptor type 1a is required for atrioventricular valve formation. Ann. Thorac. Surg. 85, 2090–8.

Kisanuki, Y. Y., Hammer, R. E., Miyazaki, J., Williams, S. C., Richardson, J. A. and Yanagisawa, M. (2001). Tie2-Cre transgenic mice: a new model for endothelial cell-lineage analysis in vivo. Dev. Biol. 230, 230–42.

Lein, E. S. Hawrylycz, M. J. Ao, N. Ayres, M. Bensinger, A. Bernard, A. Boe, A. F. Boguski, M. S. Brockway, K. S. Byrnes, E. J. et al. (2007). Genome-wide atlas of gene expression in the adult mouse brain. Nature 445, 168–76.

Lin, C. J., Lin, C. Y., Chen, C. H., Zhou, B. and Chang, C. P. (2012). Partitioning the heart: mechanisms of cardiac septation and valve development. Development 139, 3277–99.

Lopez, I., Bafalliu, J. A., Bernabe, M. C., Garcia, F., Costa, M. and Guillen-Navarro, E. (2006). Prenatal diagnosis of de novo deletions of 8p23.1 or 15q26.1 in two fetuses with diaphragmatic hernia and congenital heart defects. Prenat. Diagn. 26, 577–80.

Ma, L., Lu, M. F., Schwartz, R. J. and Martin, J. F. (2005). Bmp2 is essential for cardiac cushion epithelial-mesenchymal transition and myocardial patterning. Development 132, 5601–11.

Mandel, H., Shemer, R., Borochowitz, Z. U., Okopnik, M., Knopf, C., Indelman, M., Drugan, A., Tiosano, D., Gershoni-Baruch, R., Choder, M. et al. (2008). SERKAL syndrome: an autosomal-recessive disorder caused by a loss-of-function mutation in WNT4. Am. J. Hum. Genet. 82, 39–47.

Markwald, R. R., Krook, J. M., Kitten, G. T. and Runyan, R. B. (1981). Endocardial cushion tissue development: structural analyses on the attachment of extracellular matrix to migrating mesenchymal cell surfaces. Scan Electron Microsc. Pt.2, 261–74.

Mootha, V. K., Lindgren, C. M., Eriksson, K. F., Subramanian, A., Sihag, S., Lehar, J., Puigserver, P., Carlsson, E., Ridderstrale, M., Laurila, E. et al. (2003). PGC-1alpha-responsive genes involved in oxidative phosphorylation are coordinately downregulated in human diabetes. Nat. Genet. 34, 267–73.

Okubo, A., Miyoshi, O., Baba, K., Takagi, M., Tsukamoto, K., Kinoshita, A., Yoshiura, K., Kishino, T., Ohta, T., Niikawa, N. et al. (2004). A novel GATA4 mutation completely segregated with atrial septal defect in a large Japanese family. J. Med. Genet. 41, e97.

Pehlivan, T., Pober, B. R., Brueckner, M., Garrett, S., Slaugh, R., Van Rheeden, R., Wilson, D. B., Watson, M. S. and Hing, A. V. (1999). GATA4 haploinsufficiency in patients with interstitial deletion of chromosome region 8p23.1 and congenital heart disease. Am. J. Med. Genet. 83, 201–6.

Pendeville, H., Winandy, M., Manfroid, I., Nivelles, O., Motte, P., Pasque, V., Peers, B., Struman, I., Martial, J. A. and Voz, M. L. (2008). Zebrafish Sox7 and Sox18 function together to control arterial-venous identity. Dev. Biol. 317, 405–16.

Pierpont, M. E., Basson, C. T., Benson, D. W., Jr., Gelb, B. D., Giglia, T. M., Goldmuntz, E., McGee, G., Sable, C. A., Srivastava, D. and Webb, C. L. (2007a). Genetic basis for congenital heart defects: current knowledge: a scientific statement from the American Heart Association Congenital Cardiac Defects Committee, Council on Cardiovascular Disease in the Young: endorsed by the American Academy of Pediatrics. Circulation 115, 3015–38.

Pierpont, M. E., Basson, C. T., Benson, D. W., Jr., Gelb, B. D., Giglia, T. M., Goldmuntz, E., McGee, G., Sable, C. A., Srivastava, D., Webb, C. L. et al. (2007b). Genetic basis for congenital heart defects: current knowledge: a scientific statement from the American Heart Association Congenital Cardiac Defects Committee, Council on Cardiovascular Disease in the Young: endorsed by the American Academy of Pediatrics. Circulation 115, 3015–38.

Reamon-Buettner, S. M. and Borlak, J. (2005). GATA4 zinc finger mutations as a molecular rationale for septation defects of the human heart. J. Med. Genet. 42, e32.

Rivera-Feliciano, J. and Tabin, C. J. (2006). Bmp2 instructs cardiac progenitors to form the heart-valve-inducing field. Dev. Biol. 295, 580–8.

Sarkozy, A., Conti, E., Neri, C., D’Agostino, R., Digilio, M. C., Esposito, G., Toscano, A., Marino, B., Pizzuti, A. and Dallapiccola, B. (2005). Spectrum of atrial septal defects associated with mutations of NKX2.5 and GATA4 transcription factors. J. Med. Genet. 42, e16.

Shimokawa, O., Miyake, N., Yoshimura, T., Sosonkina, N., Harada, N., Mizuguchi, T., Kondoh, S., Kishino, T., Ohta, T., Remco, V. et al. (2005). Molecular characterization of del(8)(p23.1p23.1) in a case of congenital diaphragmatic hernia. Am. J. Med. Genet. A 136, 49–51.

Slavotinek, A., Lee, S. S., Davis, R., Shrit, A., Leppig, K. A., Rhim, J., Jasnosz, K., Albertson, D. and Pinkel, D. (2005). Fryns syndrome phenotype caused by chromosome microdeletions at 15q26.2 and 8p23.1. J. Med. Genet. 42, 730–6.

Subramanian, A., Tamayo, P., Mootha, V. K., Mukherjee, S., Ebert, B. L., Gillette, M. A., Paulovich, A., Pomeroy, S. L., Golub, T. R., Lander, E. S. et al. (2005). Gene set enrichment analysis: a knowledge-based approach for interpreting genome-wide expression profiles. Proc. Natl. Acad. Sci. U.S.A. 102, 15545–50.

Takash, W., Canizares, J., Bonneaud, N., Poulat, F., Mattei, M. G., Jay, P. and Berta, P. (2001). SOX7 transcription factor: sequence, chromosomal localisation, expression, transactivation and interference with Wnt signalling. Nucleic Acids Res. 29, 4274–83.

Tan, T. Y., Gonzaga-Jauregui, C., Bhoj, E. J., Strauss, K. A., Brigatti, K., Puffenberger, E., Li, D., Xie, L., Das, N., Skubas, I. et al. (2017). Monoallelic BMP2 Variants Predicted to Result in Haploinsufficiency Cause Craniofacial, Skeletal, and Cardiac Features Overlapping Those of 20p12 Deletions. Am. J. Hum. Genet. 101, 985–994.

Taniguchi, K., Hiraoka, Y., Ogawa, M., Sakai, Y., Kido, S. and Aiso, S. (1999). Isolation and characterization of a mouse SRY-related cDNA, mSox7. Biochim. Biophys. Acta. 1445, 225–31.

Wang, Y., Wu, B., Chamberlain, A. A., Lui, W., Koirala, P., Susztak, K., Klein, D., Taylor, V. and Zhou, B. (2013). Endocardial to myocardial notch-wnt-bmp axis regulates early heart valve development. PLoS One 8, e60244.

Wat, M. J., Beck, T. F., Hernandez-Garcia, A., Yu, Z., Veenma, D., Garcia, M., Holder, A. M., Wat, J. J., Chen, Y., Mohila, C. A. et al. (2012). Mouse model reveals the role of SOX7 in the development of congenital diaphragmatic hernia associated with recurrent deletions of 8p23.1. Hum. Mol. Genet. 21, 4115–25.

Wat, M. J., Shchelochkov, O. A., Holder, A. M., Breman, A. M., Dagli, A., Bacino, C., Scaglia, F., Zori, R. T., Cheung, S. W., Scott, D. A. et al. (2009). Chromosome 8p23.1 deletions as a cause of complex congenital heart defects and diaphragmatic hernia. Am. J. Med. Genet. A. 149A, 1661–77.

White, J. J., Arancillo, M., Stay, T. L., George-Jones, N. A., Levy, S. L., Heck, D. H. and Sillitoe, R. V. (2014). Cerebellar zonal patterning relies on Purkinje cell neurotransmission. J. Neurosci. 34, 8231–45.

Xiong, Y., Zhou, B. and Chang, C.-P. (2012). Analysis of the Endocardial-to-Mesenchymal Transformation of Heart Valve Development by Collagen Gel Culture Assay. In Cardiovascular Development: Methods and Protocols, (eds X. Peng and M. Antonyak), pp. 101–109. Totowa, NJ: Humana Press.

Yaylaoglu, M. B., Titmus, A., Visel, A., Alvarez-Bolado, G., Thaller, C. and Eichele, G. (2005). Comprehensive expression atlas of fibroblast growth factors and their receptors generated by a novel robotic in situ hybridization platform. Dev. Dyn. 234, 371–86.

Zhang, C., Basta, T. and Klymkowsky, M. W. (2005). SOX7 and SOX18 are essential for cardiogenesis in Xenopus. Dev. Dyn. 234, 878–91.

